# The LINC complex regulates Achilles tendon elastic modulus, Achilles and tail tendon collagen crimp, and Achilles and tail tendon lateral expansion during early postnatal development

**DOI:** 10.1101/2023.11.13.566892

**Authors:** Nicholas M. Pancheri, Jordan T. Daw, Destinee Ditton, Nathan R. Schiele, Scott Birks, Gunes Uzer, Calvin L. Jones, Brian T. Penney, Sophia K. Theodossiou

**Affiliations:** Chemical & Biological Engineering, University of Idaho, Moscow, ID 83844, United States; Mechanical & Biomedical Engineering, Boise State University, Boise, ID 83725, United States

**Keywords:** Tendon, development, LINC, KASH-domain, SUN-domain, nesprin, nuclear mechanosensing, mechanotransduction, mechanobiology, tenogenesis

## Abstract

There is limited understanding of how mechanical signals regulate tendon development. The nucleus has emerged as a major regulator of cellular mechanosensation, via the linker of nucleoskeleton and cytoskeleton (LINC) protein complex. Specific roles of LINC in tenogenesis have not been explored. In this study, we investigate how LINC regulates tendon development by disabling LINC-mediated mechanosensing via dominant negative (dn) expression of the Klarsicht, ANC-1, and Syne Homology (KASH) domain, which is necessary for LINC to function. We hypothesized that LINC regulates mechanotransduction in developing tendon, and that disabling LINC would impact tendon mechanical properties and structure in a mouse model of dnKASH. We used Achilles (AT) and tail (TT) tendons as representative energy-storing and limb-positioning tendons, respectively. Mechanical testing at postnatal day 10 showed that disabling the LINC complex via dnKASH significantly impacted tendon mechanical properties and cross-sectional area, and that effects differed between ATs and TTs. Collagen crimp distance was also impacted in dnKASH tendons, and was significantly decreased in ATs, and increased in TTs. Overall, we show that disruption to the LINC complex specifically impacts tendon mechanics and collagen crimp structure, with unique responses between an energy-storing and limb-positioning tendon. This suggests that nuclear mechanotransduction through LINC plays a role in regulating tendon formation during neonatal development.

## Introduction

Healthy tendon mechanical properties develop through a complex maturation process that is at least partially mediated by mechanical loading. Distinct mechanotransducive pathways result in different cellular responses, including changes to transcription and translation, collagen production, and cell proliferation, migration, and senescence^15^. The link between the actin cytoskeleton and the nucleus has emerged as a key cellular mechanosensation and mechanotransduction regulator^17,20^. A predominant mechanism through which the nucleus transduces external mechanical signals to nuclear response is the linker of nucleoskeleton and cytoskeleton (LINC) complex, which regulates cellular differentiation and has been reviewed in detail^7,11,13,42,45,56^. LINC complex and related nuclear envelope mutations are associated with number of musculoskeletal diseases^6^. Recently our group showed that disabling LINC function in pre-osteoblasts reduced bone quality, suggesting a regulatory role of LINC complex in musculoskeletal system^4^. However, LINC complex’s specific roles in regulating tendon formation are unknown. Given the high – but untested – likelihood that gene expression and phenotypical changes associated with nuclear strain transfer signals through the LINC complex impact tendon formation, there is a need to explore LINC as a regulatory mechanism during tendon formation.

The LINC complex tethers the cytoskeleton to the outer nuclear membrane (ONM), where it then spans the perinuclear space (PNS), anchors in the inner nuclear membrane (INM), and extends into the nuclear cytoskeleton to interact with multiple nuclear proteins^20^. Nesprin 1 and 2 (1/2), commonly referred to as ‘giant’ Nesprin, couple to the actin cytoskeleton at the N-terminus to receive incoming mechanical stimuli. The C-terminus of Nesprin 1/2 penetrates into the PNS, and contains a highly-conserved Klarsicht, ANC-1, and Syne Homology (KASH) domain. The KASH domain binds to the family of Sad1p, UNC-84 (SUN) proteins that span the INM and anchor to the nucleoskeletal lamin A/C system (**Figure 1**). SUN1 and 2 trimerize and bind three distinct KASH domains and are the primary mechanism through which Nesprins bind the nuclear envelope. SUN1 and 2 contribute to Nesprin binding, and singular depletion of only one of these proteins retains LINC complex efficacy^10^, suggesting potential signaling redundancy. However, SUN1 binds even in the absence of lamin A/C, whereas SUN2 appears lamin A/C dependent^12^. Further, only SUN1 directly interacts with nuclear pore complexes (NPC)^28^, which facilitate access of nuclear-translocation proteins, including β-catenin and yes-associated protein/ transcriptional coactivator with PDZ-binding motif (YAP/TAZ), into the nucleus^35^. Notably, loss of KASH functionality disables the LINC complex and associated nuclear mechanosensing.

**Figure 1.**
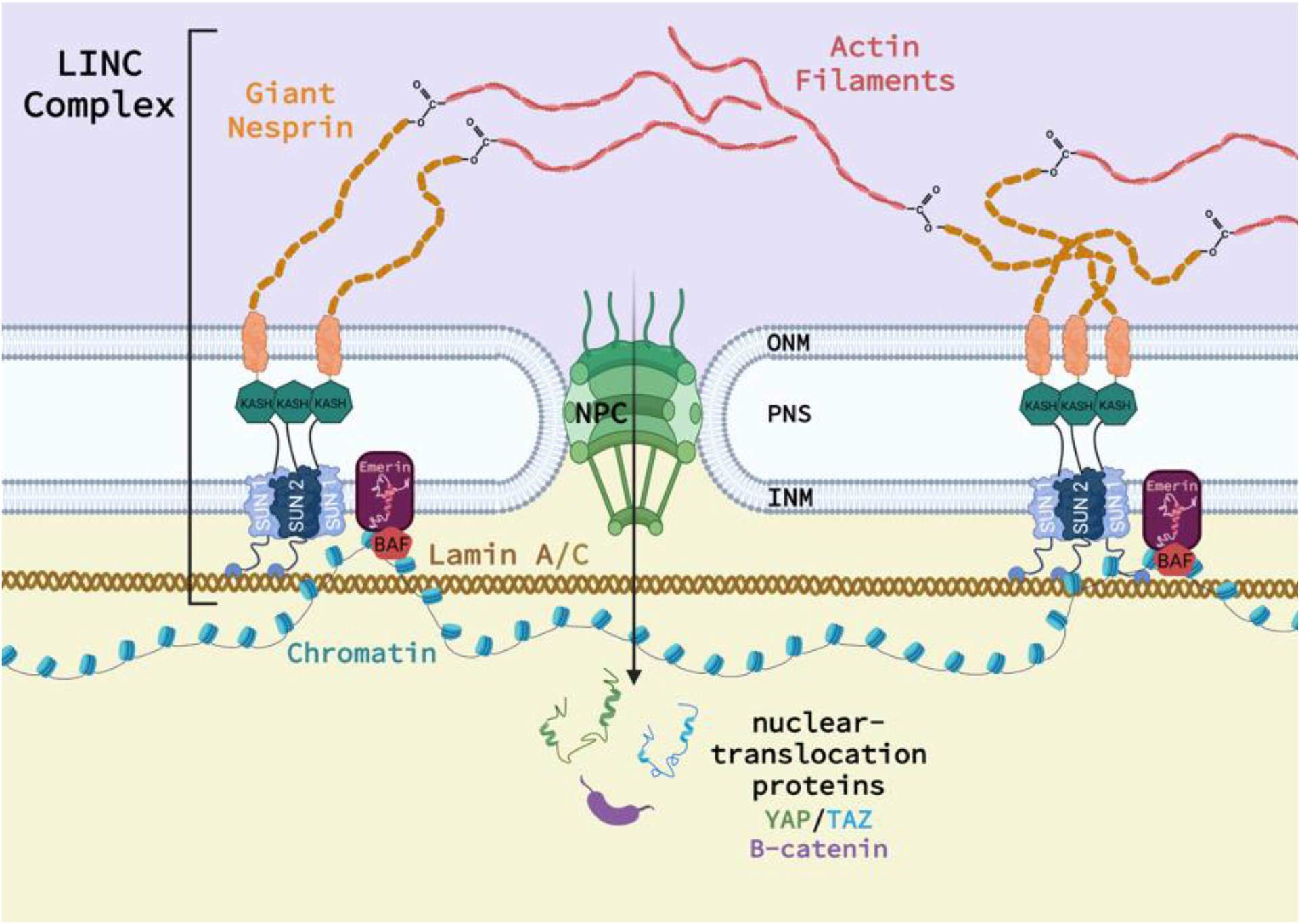
Schematic of the relationship between the LINC complex, Nesprin, Sun, KASH, and the cell nucleus. Additional proteins not investigated in this study are shown.

Little is known regarding tissue-level effects of LINC, particularly within connective tissue, though recent literature is elucidating LINC’s role in cell behavior. LINC disruption in fibroblastic cell lines impairs the nucleo-cytoskeleton’s ability to differentially regulate transcriptional activity between soft and stiff substrates, and also affects cell adhesion, wound repair, and ion signaling^1^. Small interfering RNA (siRNA) treatment and competitive SUN binding impaired low magnitude strain transduction in mouse mesenchymal stem cells (MSCs), which promoted adipogenesis through suppression of focal adhesion kinase (FAK) formation and Akt signaling^57^. Similarly, depletion of Sun 1/2 in MSCs decreased nuclear β-catenin levels, a noteworthy result given that β-catenin is a multi-role protein influencing tissue differentiation, including tenogenesis^55^. β-catenin’s role in tenogenesis is still being elucidated, however previous work shows that increased β-catenin activation follows an underloaded mechanical environment^59^, promotes ectopic ossification in tendon^59^, and suppresses expression of tenogenic markers in tendon-derived cells^21^. Taken together, existing data indicate that increased β-catenin and decreased mechanical loading disrupt normal tendon development.

It is well-established that mechanotransducive processes have critical roles in tenogenesis^19,31,58^, but many mechanotransducive tenogenic mechanisms remain enigmatic. Our work implicates Akt signaling^50^, and has summarized the role of mechanosensitive cell-cell junction proteins, particularly N-cadherin and Connexin-43^51^, on tenogenic MSCs and tendon cells, respectively. As LINC is a nuclear mechanostransducer and β-catenin/Akt modular, LINC’s potential regulatory role in tenogenesis warrants further investigation.

In this study, we disrupted the LINC complex in a murine *in vivo* model of postnatal tendon development, and assessed how LINC impacted the resulting cell and tendon tissue morphology, mechanical properties, and protein production. We hypothesized that disruption of the LINC complex would impair tendon structural and mechanical properties via alterations to the underlying collagen structure and dysregulation of tissue growth, with specific changes dependent upon tendon type (i.e., energy-storing vs positional tendon) during postnatal development. To test this, we bred mice with a dominant negative KASH domain (dnKASH) targeted to scleraxis expressing cells (i.e., tendon cells)^41^ and harvested the Achilles tendons (AT), as a representative energy-storing tendon, and tail tendons (TT), as a representative positional tendon, at postnatal day(P)10. We assessed tendon mechanical properties, tissue and cellular morphology, and collagen content and morphology. Our results provide a novel perspective of the potential role of nuclear mechanosensing in regulating tendon structure and function during development.

## Methods and Materials

### Animals and LINC Disabling via dnKASH induction

Scleraxis green fluorescent protein (Scx-GFP) mice^41^ were generously provided by Dr. Ronen Schweitzer. Mouse strains included Tg(CAG-LacZ/EGFP-KASH2) aka KASH2 described previously^43,44^. We have recently utilized a similar strategy using other bone specific cre models^4,5^. In short, hemizygous ScxGFP mice were crossed with floxed KASH2 mice to generate a ScxGFP-LINC disrupted (++) murine model. Briefly, the LINC disruption mechanism is initiated upon cre recombination: a LacZ containing lox-stop-lox cassette upstream of an eGFP-KASH2 ORF is excised allowing for the overexpression an eGFP-KASH2 fusion protein which saturates available Sun/KASH binding in a dominant-negative manner. Genotyping was performed on tissue biopsies via real-time PCR probing for LacZ and Cre (Transnetyx, Cordova, TN) to determine experimental animals and controls. Cre(+)/LacZ(+) animals were considered experimental and Cre(+)/LacZ(-) mice were used as controls. Mice were not treated with tetracycline or its derivatives as Scx-Cre activation did not cause significant fetal lethality. Pups were housed in individually ventilated cages with the dams under controlled conditions until sacrifice. Ad libitum access to food and water was allowed. All procedures were approved by the Boise State University Institutional Animal Care and Use Committee.

### Achilles and Tail Tendon Mechanical Evaluation

Mice were sacrificed at postnatal day(P)10 using previously established methods. P10 represents relatively early stages of postnatal tendon formation and when full weight bearing locomotor behavior has initially developed in mice and rats^2,49^. Achilles tendons (AT) (n=10-13/treatment) and tail tendons (TT) (n=12-13/treatment) were mechanically evaluated via a custom small-scale tensile load frame as previously described^49,52^. Briefly, the skin was removed from tails, and the proximal end of the tail was gripped while each TT was gently teased away from the tail using ultra fine dissection tweezers. Similarly, skin was removed from the rear hind limb and then the AT was isolated by severing through the quadriceps muscle proximally and the calcaneal bone distally. The residual muscle of the myotendinous junction (MTJ) and calcaneal bone was mounted in the upper and lower grips, respectively. The MTJ was secured to the upper clamp using rubber sheeting and cyanoacrylate, and the calcaneal bone was secured using fine grit sandpaper, cyanoacrylate, and rubber sheeting. TTs were adhered to cardboard c-clamps and mounted in the upper and lower grips. Tendon cross-sectional area and gauge length were measured from images acquired with a digital camera (Thorcam DCC1645C, Thorlabs INC., Newton, NJ). TTs were assumed to have a circular cross section and ATs an elliptical cross section, based upon previous literature^8,25,37^. Tendons were kept hydrated with Dulbecco’s phosphate buffered saline (DPBS, Gibco, White Plains, NY) for the duration of testing. Preconditioning was applied to ATs with 10 cycles of 1% strain s^-1^ and immediately followed by pull to failure at 0.1 mm/s^49^. TTs were not preconditioned due to the fragility of the P10 TTs, but slack was carefully removed from each sample via application of a 0.1 N preload prior to pull-to-failure tensile testing. A 150 g capacity load cell (Honeywell, Columbus, OH) measured forces, and a custom LabVIEW (National Instruments, Austin, TX) program recorded force and displacement data. Recorded force, displacement, and cross-sectional area were then used to determine maximum force, displacement at maximum force, maximum stress, and strain. Stiffness and elastic modulus were derived from the linear regions (where the slope of a line of fit had an R^2^ > 0.9, average R^2^ =0.96) of the force-displacement and stress-strain curves, respectively.

### SHG and SEM Imaging to Assess Collagen Morphology and Crimp

Collagen morphology and crimp structure were evaluated using two distinct imaging modalities (second harmonic generation or scanning electron microscopy). In both methods, tendons were isolated via similar dissection methods as described above. ATs (n=3) or TTs (n=4) were excised from the surrounding tissue and immediately fixed in a tension-free state using 10% formalin (for SHG) or 4% *para-*formaldehyde (for SEM) (Sigma), and stored at 4 °C.

#### Second Harmonic Generation (SHG) Microscopy

For ATs (n=3 dnKASH), the myotendionous junction and calcaneus were removed following fixation and placed onto a glass slide and saturated with PBS. TTs (n=5 dnKASH) were placed directly onto the glass slide with PBS. For each sample, images were acquired in the center, left, and right positions of the tissue using an Olympus Fluoview 1000 Confocal Microscope (Olympus). Based upon previously established methods^52^, ImageJ was used to directly measure the crest-to-crest distances of 5 distinct crimp fibrils within each image. The mean crimp distance for each image was then averaged with the remaining two image locations to calculate an average measurement for each sample.

#### Scanning Electron Microscopy (SEM)

ATs (n=5 control, 8 dnKASH) and TTs (n=5 control, 6 dnKASH) were prepared for SEM and imaged the day after the dehydration procedures concluded. Tendons were dehydrated in increasing concentrations of ethanol following a standard protocol for cellular sample SEM preparation^22^. Dehydrated samples were sectioned with a razor blade, mounted onto SEM sample studs using adhesive carbon tabs, and sputter-coated with gold particles for 30 seconds using an Emitech Mini Sputter Coater (SC7620, Quorum Technologies). Samples were imaged using a Zeiss EVO MA10 SEM (Zeiss), with the electron beam set to 5kV.

### Fluorescence Imaging to Assess Whole Tendon Cellularity

To evaluate P10 tendon cellularity, ATs and TTs were fixed in 4% PFA and stained with 4′,6- diamidino-2-phenylindole (DAPI) (Invitrogen) to identify cell nuclei. Stained tendons were imaged at 10X on an Olympus ix70 inverted fluorescence microscope (Olympus, New York, NY). The presence of scleraxis-positive cells was also confirmed this way. FIJI (ImageJ) was combined with a custom ImageJ program to separate and count cell nuclei in the tendon samples based on shape and contrast. Average cell nuclei counts were averaged for n=3 images per tendon type and condition (control or dnKASH). The results of the program were validated against manual counts obtained by an independent observer, and no significant differences were detected.

### Statistical Analyses

Mechanical testing results and crimp measurements were evaluated using nonparametric unpaired, two-tailed Mann-Whitney U tests (Prism 9, Graphphad, LaJolla, CA). Mann-Whitney U tests were selected over standard unpaired t-tests because of similarly skewed distributions between groups, rather than a normal distribution. Significance was set as p<0.05, with results reported as mean ± standard deviation.

## 5. Results

### Disabling LINC via dnKASH Expression Decreases Achilles Tendon Elastic Modulus and Maximum Stress, and Increases Cross-Sectional Area

Descriptive statistics of AT and TT mechanical properties and gauge lengths are shown in Table 1. Representative force-displacement and stress-strain curves for dnKASH (++) and control (+-) demonstrating the linear region used to calculate P10 AT mechanical properties (Supplemental Figure 1A, B) following LINC disabling. AT elastic modulus decreased 5.5-fold (p=0.0065) and maximum stress decreased by 3-fold (p=0.048) in dnKASH (++) samples, compared to tendons from mice without the dnKASH (+-) (Figure 2E, G). AT cross-sectional area increased (p=0.0089) with the dnKASH (Figure C). Maximum force (p=0.6665), displacement at maximum force (0.8859), stiffness (p=0.4366), and strain at maximum stress (p=0.7844) remained consistent between dnKASH and controls (Figure 2A, B, D, F). Overall, at P10 dnKASH significantly impacted AT material properties. Despite the increase in cross-sectional area, cell density did not differ significantly between control and dnKASH ATs (Fig. 6A, B, E).

**Figure 2.**
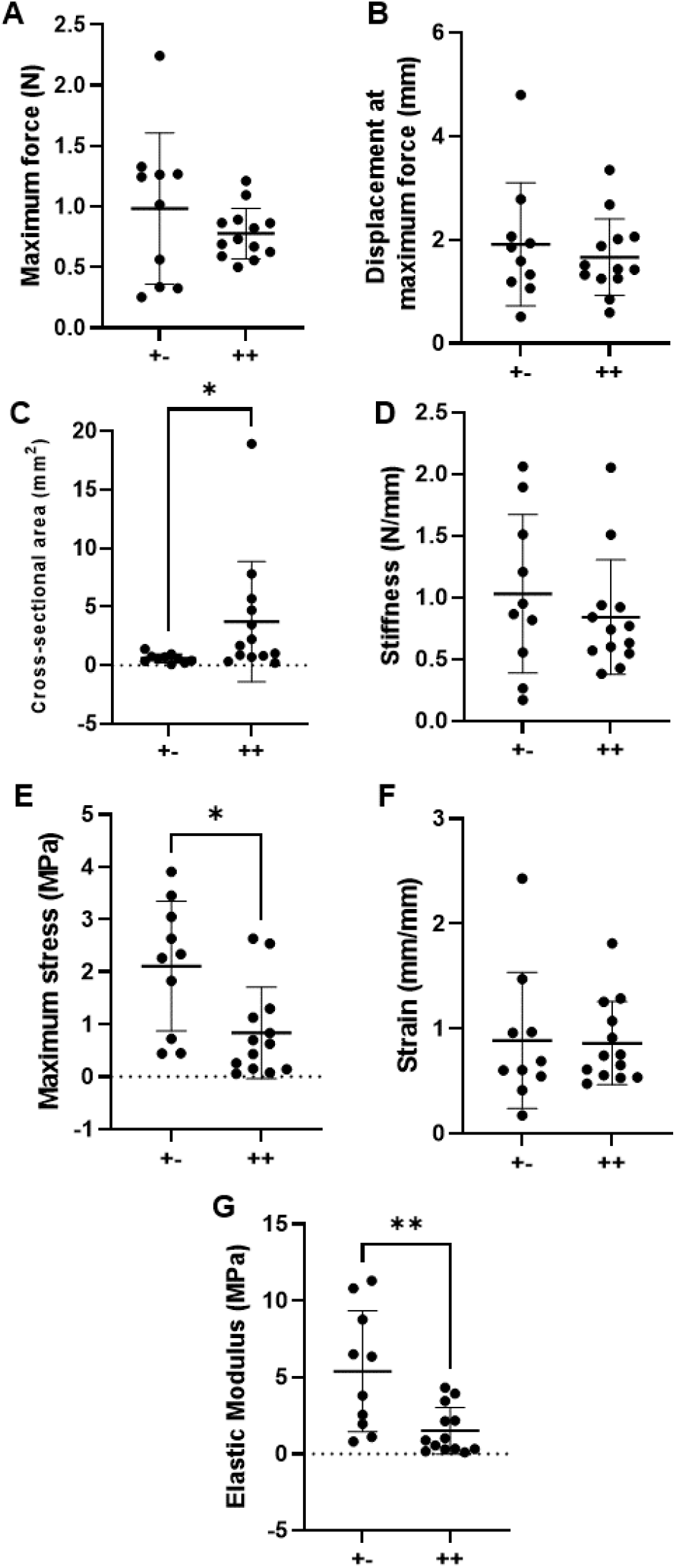
P10 Achilles tendon mechanical properties: (A) Maximum force, (B) displacement at maximum force, (C) cross-sectional area, (D) stiffness, (E) maximum stress, (F) strain, and (G) elastic modulus. dnKASH ++ ATs had a decreased (E) maximum stress and (G) elastic modulus but (C) an increased cross-sectional area, compared to +- controls. Bridging lines and asterisks denotes significance (p<0.05). Bars represent mean ± standard deviation

### dnKASH Increases Tail Tendon Maximum Stress and Decreases Cross-Sectional Area

Representative force-displacement and stress-strain graphs included (Supplemental figure 1 C, D). TT cross-sectional area decreased (p=0.0135) with dnKASH, compared to control groups (Figure 3C). In contrast to the material property changes observed in the ATs, TTs from dnKASH mice trended towards an increase in maximum tissue stress (p=0.0976) (Figure 3E). Maximum force (p=0.6404), displacement at maximum force (0.6041), stiffness (p=0.9787), elastic modulus (p=0.1225), and strain at maximum stress (p=0.3551) did not differ significantly between dnKASH and control groups (Figure 3A, B, D, F, G). Additionally, no significant differences in cell number were identified between control and dnKASH TTs (Figure 6C, D, F). Overall, dnKASH expression appeared to impact the positional TTs less than the energy-storing ATs, suggesting that differing mechanotransducive mechanisms drive positional and energy-storing tendon development.

**Figure 3.**
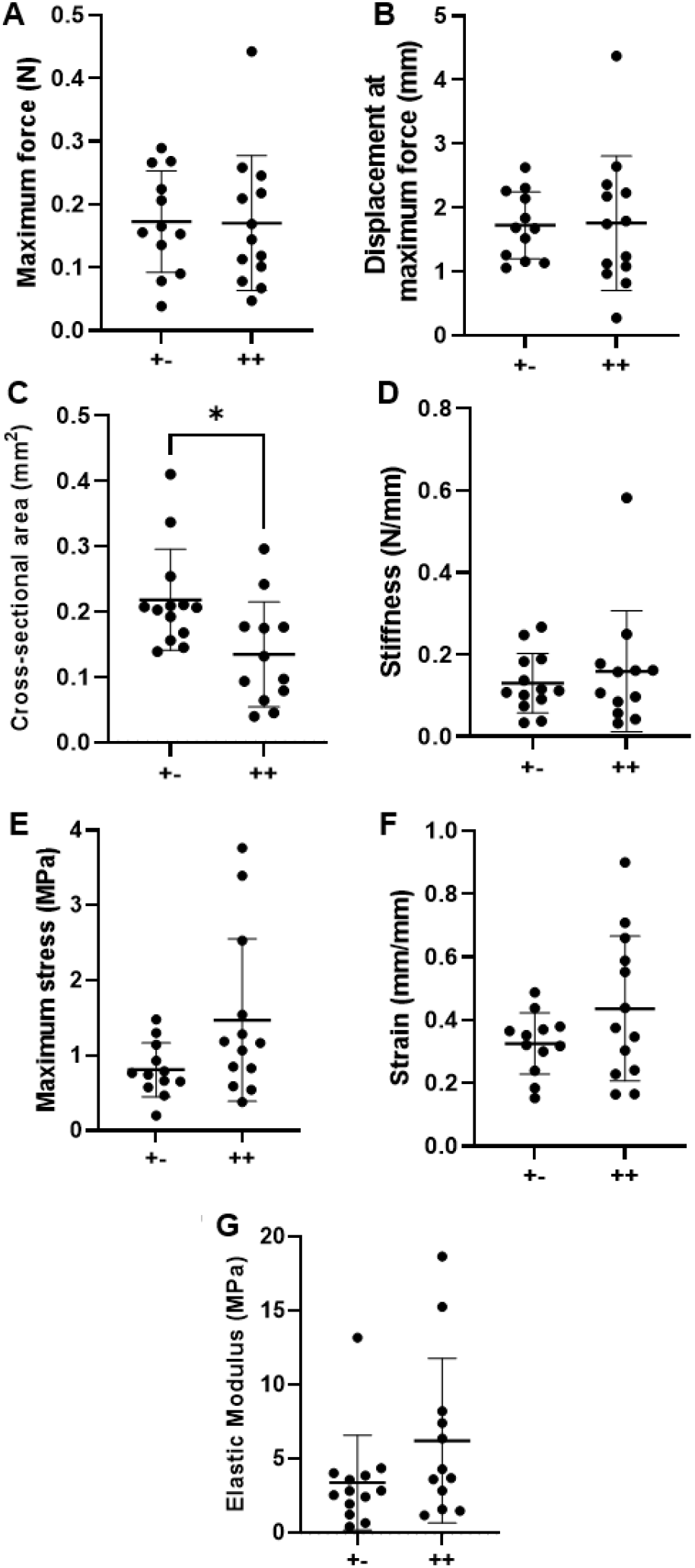
P10 tail tendon mechanical properties: (A) Maximum force, (B) displacement at maximum force, (C) cross-sectional area, (D) stiffness, (E) maximum stress, (F) strain, and (G) elastic modulus. dnKASH ++ TTs had a decreased (C) cross-sectional area, compared to +- controls. Other mechanical properties did not differ significantly between dnKASH and control TTs. Bridging lines and asterisks denotes significance (p<0.05). Bars represent mean ± standard deviation.

### dnKASH Significantly Affects Crimp Distance in Both ATs and TTs

Crimp distance was measured via SHG and SEM imaging, with no statistical differences in crimp measurements between imaging modalities when comparing with standard T-tests (Figure 4 and Supplemental Figure 2). A change in core facility equipment during the study necessitated pooling crimp distance measurements acquired from both SHG and SEM imaging to achieve a sample size of at least 5 per tendon type and condition (n=8/treatment for ATs, n=12/treatment for TTs). When comparing dnKASH, to control tendons, the crimp distance was significantly decreased in dnKASH ATs (Figure 5A) and significantly increased in dnKASH TTs (Figure 5B). There were significant differences between ATs and TT crimp distance (data not shown), as seen previously.^52^

**Figure 4.**
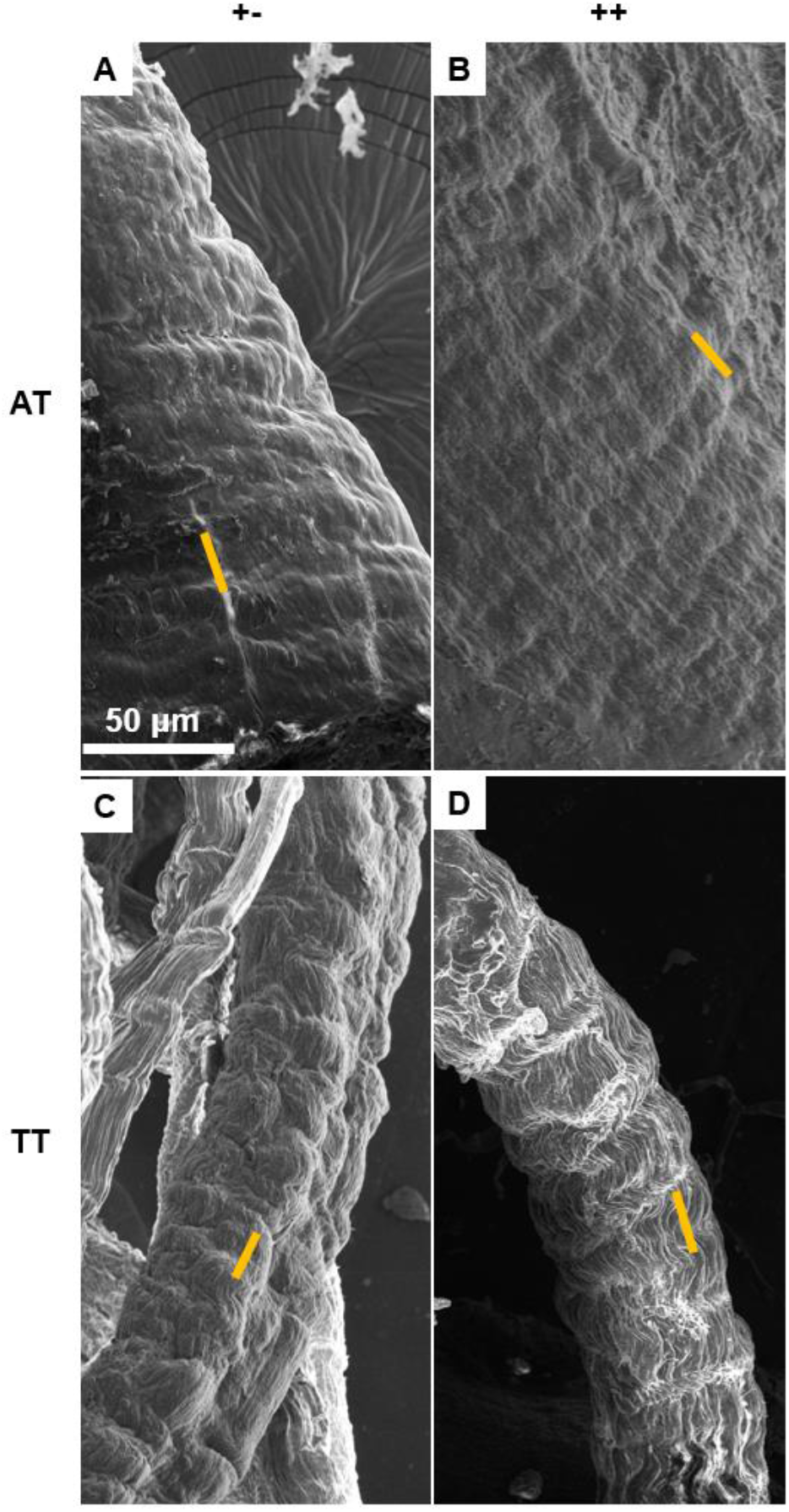
SEM micrographs of (A) control +- and (B) dnKASH ++ ATs, and (C) control +- and (D) dnKASH ++ TTs showing crimp pattern and morphology. Though crimp distances differed significantly between control and dnKASH tendons, no obvious ultrastructural or morphological differences are observed. Yellow lines show representative crimp measurements taken between crimp wave “peaks.” Crimp was measured identically in SHG and SEM images. All images captured at 800x.

**Figure 5.**
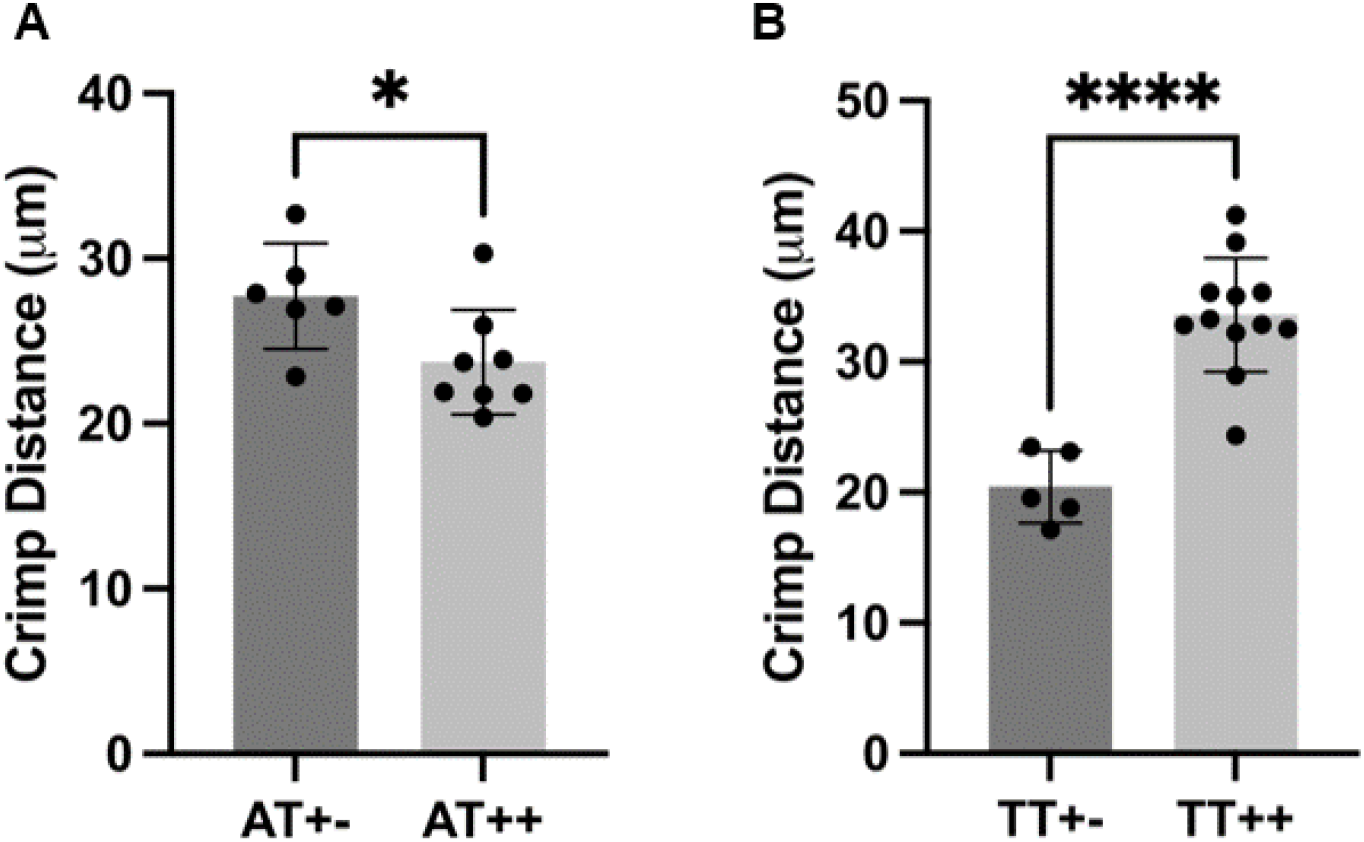
Disabling nuclear mechanosensing via dnKASH significantly impacted crimp distance in ATs and TTs. (A) dnKASH ++ ATs had significantly decreased crimp distance compared to control +- ATs. Conversely, (B) dnKASH ++ TTs had significantly increased crimp distance compared to control +- TTs.

### No Significant Differences Identified in Gross Morphology or Cellularity Between dnKASH and Controls

SEM and fluorescence microscopy were used to investigate potential changes to collagen or cellular morphology. We did not observe qualitative changes to the ultrastructural characteristics of AT or TT collagen fibrils. Fluorescence imaging confirmed GFP localization to scleraxis expressing cells, with an obvious green diffusion throughout the tissue (Supplemental Figure 4) but counting cell nuclei numbers showed no differences in total cell count (Figure 6), or cellular morphology between ATs and TTs from dnKASH mice versus controls.

**Figure 6.**
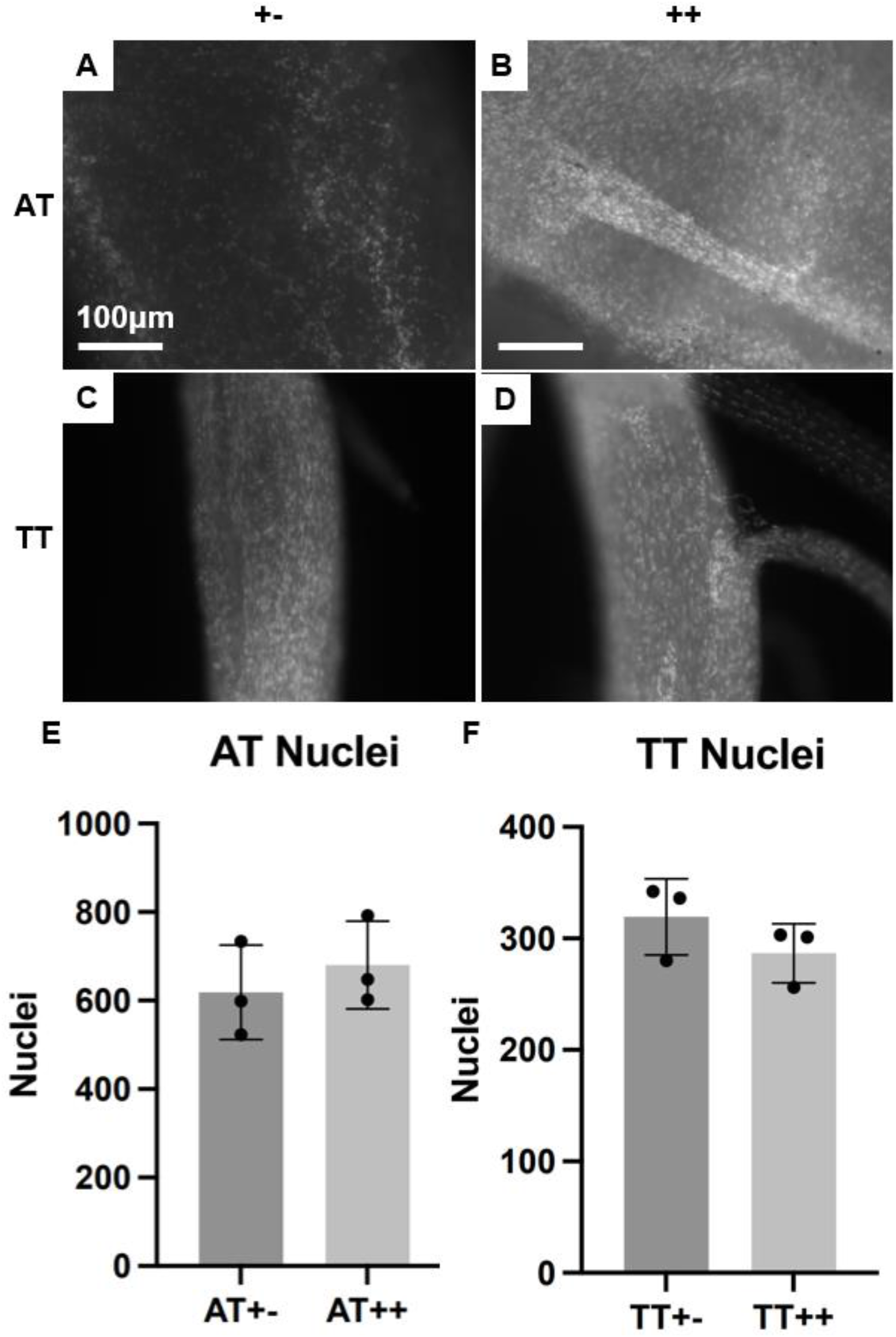
(A-D) SHG micrographs of DAPI-stained nuclei in whole (A) control +- ATs, (B) dnKASH ++ ATs, (C) control +- TTs, and (D) dnKASH ++ TTs. (E, F) Counting the nuclei showed no significant differences between dnKASH and controls for either tendon type. All scale bars as shown. All images at 10x, scale bars = 100μm.

### dnKASH Significantly Affects Tendon Marker Production in Mesenchymal Stem Cells (MSCs) (Supplemental Data)

Western Blot and phase contract imaging methods and data are presented in the supplemental section and figures. Our preliminary analysis suggests that disabling LINC via dnKASH inhibits tenogenesis and cell survival during TGFβ2-induced tendon differentiation in MSC (Supplemental Figure 3 A-L) and disrupts tenogenic marker production (Supplemental Figure M). Though the cellular pathways that support the regulatory role of LINC during tendon development will be the subject of future investigations, we are including these preliminary data as supporting evidence that LINC merits further investigation in the context of postnatal tendon development.

## Discussion

While it is established that mechanical loading affects tendon development^3,23,30,38,40,49,52^, the exact signaling mechanisms remain unclear. The primary transducer between the actin cytoskeleton and nuclear architecture is the LINC complex, which has been implicated in many cellular functions, including differentiation^42,45^, nuclear morphology^32^, and DNA repair^27,60^. While the LINC complex has been studied in recent years, little work has explored LINC’s role in tendon development. As such, the aim of this study was to exploit dnKASH mice to investigate the effects of LINC on tendon mechanical functionality during early postnatal development by evaluating mechanical properties and tissue morphology.

A dnKASH was targeted to scleraxis expressing cells (i.e., tendon cells) in mice to impair the LINC complex of developing tendons at P10. We observed the energy-storing ATs to be more severely impacted by the dnKASH than positional TTs, although TTs were also affected (Figures 2, 3). AT elastic modulus and maximum tissue stress decreased, whereas cross-sectional area increased. Together, these changes indicate a disruption in tendon development, further implicating mechanical loading as a regulator of lateral expansion in developing ATs^49^. . Other mechanical properties (maximum force, displacement at maximum force, stiffness, and strain at maximum stress) were not significantly affected. The overall collagen structure, morphology, and organization were also unaffected by the dnKASH (Figure 4A-D), suggesting a cellular and/or molecular mechanism is responsible for the observed changes in AT material properties. Unexpectedly, disabling nuclear mechanosensing affected collagen crimp distance, with the dnKASH corresponding to significantly shorter crimp distances in ATs, and significantly longer crimp distances in TTs (Figure 5). Unknown mechanisms drive these changes, but changes could be attributed to a variety of factors such as tenocyte contraction or age-related alterations in crimp distance^14,26^ . The differential effect of the dnKASH on an energy-storing versus positional tendon further highlights the unique developmental program between tendon types that is possibly driven by differences in their mechanical loading.

Ours and others’ previous work shows increased AT elastic modulus when mechanical loading is disrupted through a spinal cord transection or Botox treatment in developing rats, though these studies do not identify significant changes in collagen crimp distance^18,52^. Prior work has also shown that crimp angle decreases with age, potentially in response to stiffening of the surrounding matrix .^26^ . While crimp angle was not measured in this study, smaller crimp angles correspond to larger crimp distances, which suggests larger crimp distances may be found in relatively older tendons. The significantly smaller crimp distance measured in the dnKASH ATs may be due to delayed tendon maturation following impaired mechanosensing. Future studies will include additional imaging modalities and measure crimp angle, as both crimp distance and crimp angle may contribute to the sliding and stretching of collagen fibers, the primary tendon loading mechanism during normal physiological movement ^38,46,48^. *In vivo* models have established that alterations to the developmental loading environment significantly impact tendon mechanical properties ^18,36,40,52^. However, the specific cellular mechanisms that translate mechanical stimuli to cellular and tissue responses remain unclear.

Interestingly, the decreases in stress and elastic modulus in the ATs alongside unchanged structural properties (maximum force, displacement at maximum force, stiffness), suggest LINC signaling may affect collagen deformation mechanisms (e.g., crimp length, crimp angle, collagen crosslinking). Indeed, previous work found that chemically induced paralysis decreased embryonic chick tendon elastic modulus, which corresponded to decreased lysyl oxidase (LOX) activity^36^. LOX is the primary collagen crosslinking enzyme produced by cells, and plays a role in regulating developing tendon mechanical properties^33^. Limited mechanotransduction through LINC in tendon cells may have a similar outcome as paralysis, and impact collagen crosslinking. The LINC complex has also been implicated as a potential regulator of membrane type 1-matrix metalloproteinase (MT1-MMP), which facilitates cell migration through collagen matrix^16^. Taken together, disrupting LINC may have altered these tissue modifying enzymes (i.e., LOX, MMPs), which contributed to the observed mechanical and crimp changes. Follow-up studies will investigate how disrupting LINC-mediated mechanotransduction impacts ECM assembly and composition as well as LOX and MMP production.

Further research is needed to identify the molecular and cellular mechanisms behind the observed opposing changes in AT (increased) and TT (decreased) cross-sectional area with dnKASH. Existing literature has identified a proportional relationship between levels of lamin-A, nuclear stiffness, and collagen type I in soft and hard tissues, including cartilage and bone, with lower lamin-A levels favoring nuclear remodeling^47^. To our knowledge, no studies have compared nuclear lamin-A differences between developing ATs and TTs. However, evidence generally points to increased tissue elastic modulus corresponding with increased lamin-A content^47^. Differences in ATs and TTs may therefore be explained by AT elastic modulus corresponding to increased lamin-A content, such that perturbation of the laminar network via the dnKASH more severely impacted AT mechanical properties, compared to TTs. Conversely, lower lamin-A levels in TTs and less exposure to mechanical loading may have further blunted the impacts of dnKASH on TT mechanical properties. Mechanical changes associated with a dysregulated lamin in ATs may be driven by the increased loading ATs experience as an energy-storing tendon, versus positional TTs, resulting in alterations to lamin-dependent gene expression. LINC disruption and the associated effects on the nuclear lamina had unknown effects on tendon formation in this study, warranting further investigation.

Other factors include developmental differences between ATs and TTs. In rats, ATs had significant lateral expansion of the cross-sectional area from P1 to P10 that corresponds to increases in locomotor-related loading^49^. However, across similar timepoints, the CSA of rat TTs did not appear to increase. While we did not assess locomotor development in this study, a similar phenomenon may have occurred in these mice, affecting tendon cellularity and crimp distance. The dnKASH appeared to impact AT mechanical properties more severely than TTs. A developmentally -dependent response to the dnKASH may suggest the LINC complex also serves as a type of “mechanostat” that helps regulate tissue response to loading during maturation, and could influence cell growth and differentiation during development.

This study has some limitations. There are significant challenges associated with isolating and mounting neonatal mouse tendon tissue for mechanical testing, notably the small sample sizes, fragility, and potential damage during tissue dissection. Testing errors were minimized by strict exclusion of mechanical testing data that showed slipping. Data were also excluded following tissue damaged during mounting or dissection, or contact with the cyanoacrylate glue. We previously complied a detailed summary of neonatal testing limitations^49^. Other limitations may include the potency and accuracy of the scleraxis-linked dnKASH. Relative fold changes comparing reduction in KASH activity with the dnKASH, relative to the scleraxis-linked controls alone, were not assessed in this study, but have been published elsewhere^41^. GFP tag accuracy to scleraxis expressing cells was qualitatively verified using fluorescence imaging but was not quantitively examined. Finally, sex was not controlled for during sample testing, although sex-related differences (e.g., body mass, collagen content, mechanical properties, etc.) do not occur until after 4 weeks in mice^34^. Lastly, though it is not ideal to pool SEM and SHG crimp measurements, the measured distances were not statistically significant (Supplemental Figure 2).

The results of this study begin to untangle the potential regulatory role of LINC in postnatal tendon development. LINC appears to regulate lateral expansion and collagen crimp distance in both energy-storing ATs and positional TTs, although the extend of LINC regulation may be dependent upon tendon type. Our findings emphasize the need for further investigation into the molecular and cellular mechanisms that underlie mechanostransduction during early tendon development. Future work will explore the genetic, cellular and molecular (i.e. collagen cross-linking) changes in tendon resulting from LINC disruption to identify additional regulatory mechanisms, as well as the emerging differences in energy-storing and positional tendon development. These results are an important step towards defining the role of nuclear mechanosensing on tendon formation and mechanical functionality.

## Acknowledgements

The authors thank Dr. Ronen Schweitzer for providing the Scleraxis-gFP mice. This project was made possible by a NASA EPSCoR Research Initiation Grant, and Beckman Scholars Award from the Arnold and Mabel Beckman Foundation (to NMP). We acknowledge support from the Institutional Development Awards (IDeA) from the National Institute of General Medical Sciences of the National Institutes of Health under Grants #P20GM103408, P20GM109095 P20GM109095, AG059923, AR075803, and 1C06RR020533, and from the National Science Foundation under Grants 1929188 and 2025505. We also acknowledge support from The Biomolecular Research Center at Boise State, BSU-Biomolecular Research Center, RRID:SCR_019174, with funding from the National Science Foundation, Grants #0619793 and #0923535; the M. J. Murdock Charitable Trust; Lori and Duane Stueckle, and the Idaho State Board of Education.

## Author Contributions Statement

All authors have read and approved the final submitted manuscript.

Nicholas M. Pancheri: concept/design, data analysis/interpretation, manuscript writing. Jordan T. Daw: data analysis/interpretation, manuscript writing.

Destinee Ditton: data analysis/interpretation, manuscript writing.

Nathan R. Schiele: concept/design, data analysis/interpretation, financial support, manuscript writing.

Scott Birks: data analysis/interpretation.

Gunes Uzer: concept/design, data analysis/interpretation, financial support, manuscript writing. Calvin L. Jones: data analysis/interpretation, manuscript writing.

Brian T. Penney: data analysis/interpretation, manuscript writing.

Sophia K. Theodossiou: concept/design, data analysis/interpretation, financial support, manuscript writing, final approval of manuscript..

