## Supplemental Data and Figures for "The LINC complex regulates Achilles tendon elastic modulus, Achilles and tail tendon collagen crimp, and Achilles and tail tendon lateral expansion during early postnatal development"

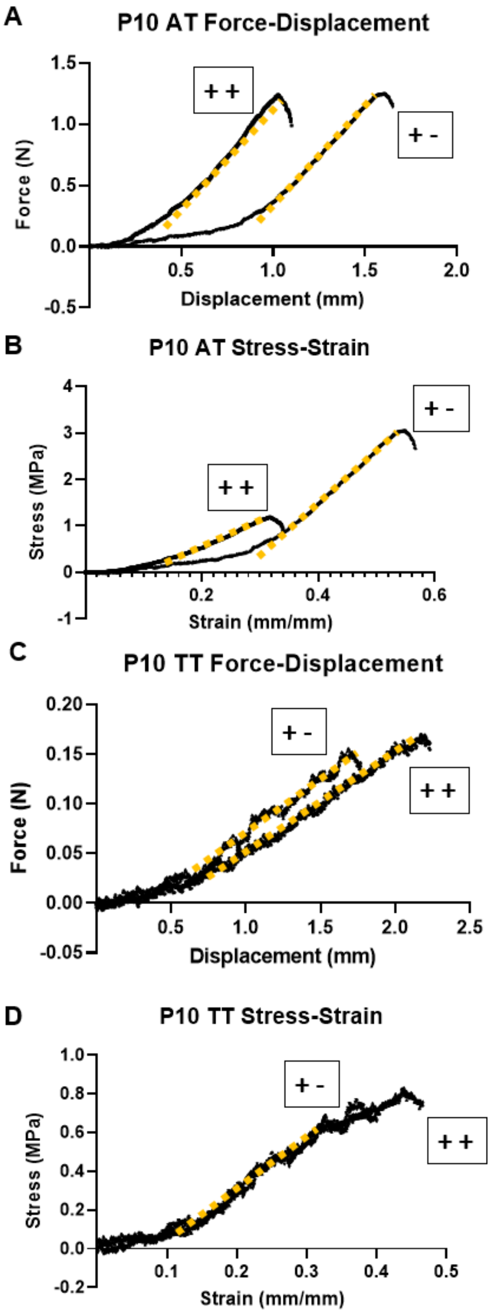


Supplemental figure 1. Representative force-displacement and stress-strain curves for (A,B) ATs and (C,D) TTs. Yellow dashed lines show the linear region used to calculate stiffness and elastic modulus. All yellow lines have a fit with the $R^2$ value ≥ 0.95; median $R^2$ = 0.998.

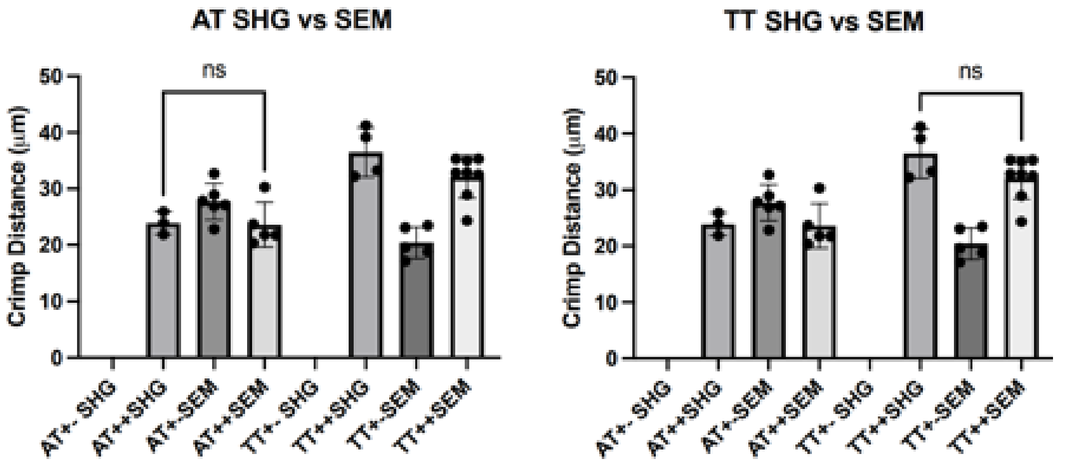


Supplemental Figure 2. Crimp distance did not differ significantly between ATs and TTs imaged via second harmonic generation (SHG) and scanning electron microscopy (SEM) imaging. No control +- ATs and TTs were imaged on the SHG, so no comparison is provided.

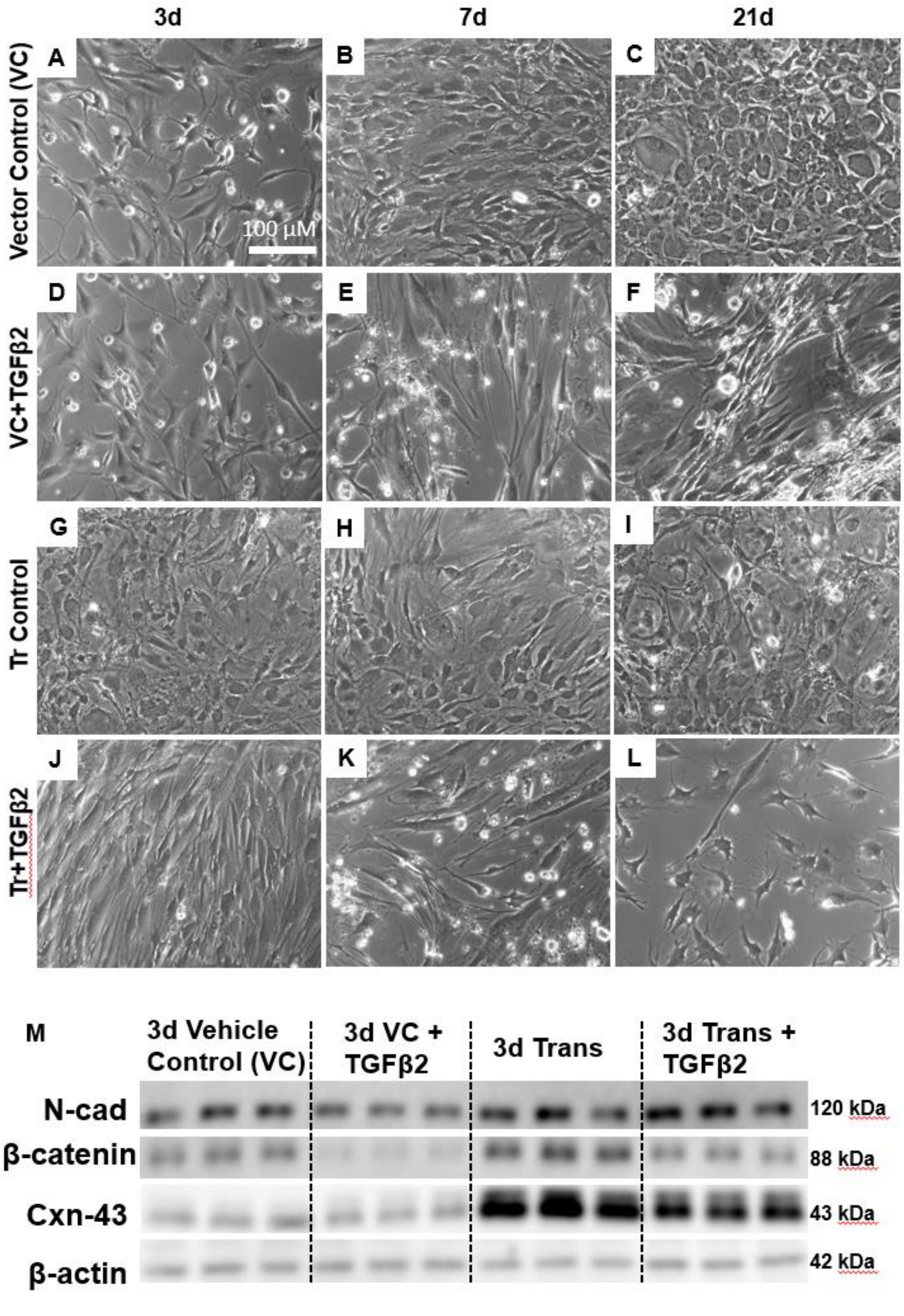


Supplemental Figure 3. Phase-contrast images (A-L) show normal mouse MSC controls (A-C) and TGFβ2-treated (D-F) cells. As shown previously[53], TGFβ2 treatment caused cells to elongate and proliferate. Transfected dnKASH cells used as controls (G-I) showed sustained growth through 21 days, but with some cell death and changes in morphology. Transfected cells treated with TGFβ2 (J-K) grew poorly, and not all replicates survived to 21 days. (M) Western Blots show that transfected MSCs do not display the previously characterized drop in N-cadherin production when treated with TGFβ2, unlike control mouse MSCs. Transfected cells appear to produce much higher levels of Connexin-43 and β-catenin, with and without TGFβ2 treatment. All images taken at 10x. Scale bar = 100 µm.

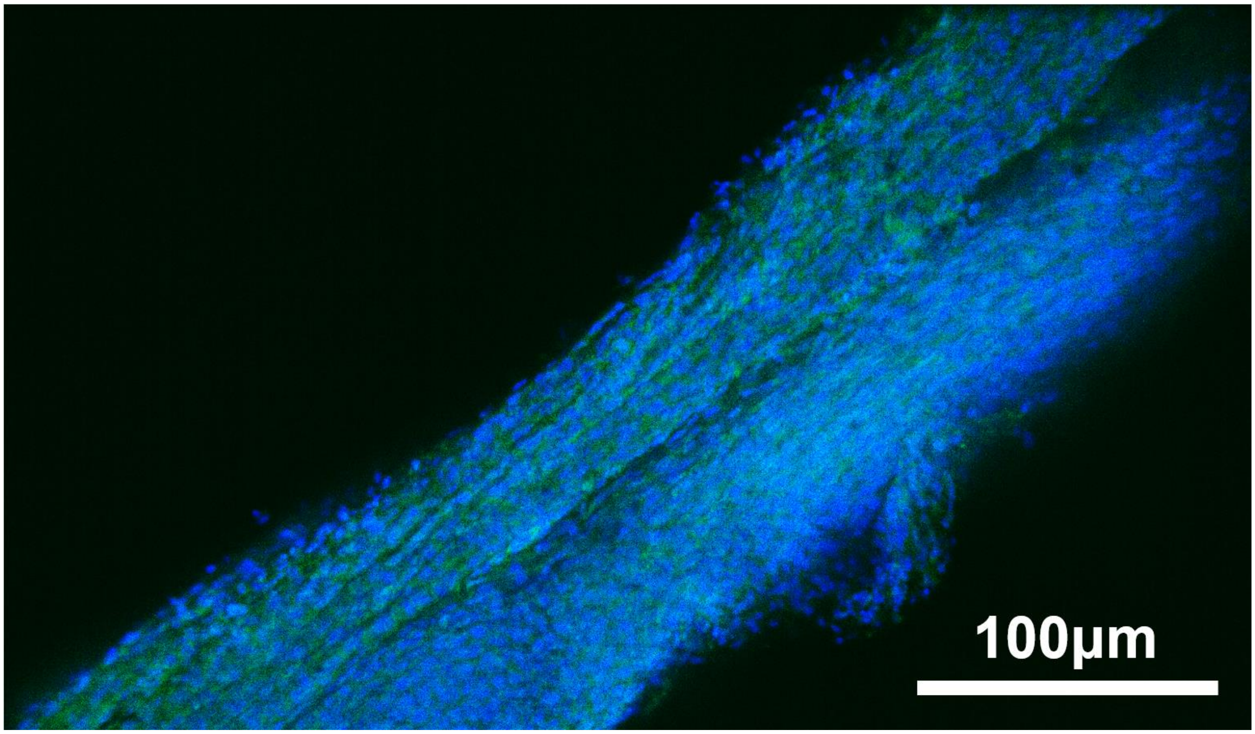


Supplemental Figure 4. Fluorescent image of whole P10 Scx-GFP ++ mouse Achilles tendon. Scleraxis expression is observed throughout the tendon, and high cellularity corresponds to the early developmental stage. Visible nuclei were used for counting. Blue = DAPI nuclear stain, green = GFP. Image taken at 10x. Scale bar = 100 μm

**Supplemental Data**

*MSC dnKASH Model*

We isolated primary mouse mesenchymal stem cells (MSCs) from multiple 8 to 10 week old B6 male donors. MSCs were extracted from 3-5 mice and the cells pooled together, following an established protocol[54,55,57]. Following the extraction protocol[39] we tested the MSCs for adipogenic and osteogenic potential using standardized methods[9], and subsequently froze the cells. MSCs were used between passages 6 and 9. To disable LINC function *in vitro* we used previously established protocols[57].Briefly, pCDH-EF1-MCS1-puro-mCherry (mCherry control) and pCDH-EF1-MCS1-puro-mCherry-Nesprin-1αKASH (dnKASH) plasmids were kindly provided by Dr. Lammerding[24]. mMSCs were transfected using 1μg DNA per 100,000 cells using LipoD293 transfection reagent (SignaGen Laboratories, Rockville, MD) according to manufacturer's instructions. 72h after the initial transfection, stably transfected cells were selected using 2μg/ml puromycin. Puromycin was used throughout the cell culture experiments to select for transfected cells.

As with the Cre animal model used in this study, transfected MSCs expressed a dominant negative nesprin-KASH (dnKASH) domain that competitively binds to SUN protein instead of endogenous nesprin, thus disabling LINC function by separating nesprin-SUN connectivity[29]. Puromycin was added to the culture media at a concentration of 2 μg/mL to select for dnKASH MSCs. All culture experiments were performed in triplicate, though some of the included supplemental data (Western Blot assays and cell morphology phase contrast imaging) was generated from preliminary experiments performed in duplicate for certain conditions; this has been noted where applicable.

MSC culture was performed using established lab protocols published elsewhere[53]. Briefly, dnKASH transfected cells were cultured in T-75 flasks (Corning) in standard growth medium (Dulbecco's Modified Eagle's Medium (DMEM) 10% fetal bovine serum (FBS), and 1% Penicillin/Streptomycin; all from Gibco) and 2μg/ml puromycin to select for plasmid-containing cells, to 70% confluency, and used between passages 6 and 9. Following standard cell passaging, the cells were detached using 0.1% trypsin (Gibco) and 5,000 cells/cm$^2$ were plated into tissue culture polystyrene (TCS) 12-well plates (Corning) and allowed to attach and equilibrate for 24 hrs. The medium was then changed to low-serum DMEM (standard DMEM supplemented with 1% FBS, 2μg/ml puromycin, and 1% Penicillin/Streptomycin) for an additional 24 hrs. Following this equilibration period, the cells were washed in warm PBS and the medium was changed to low-serum DMEM and 2μg/ml puromycin, or low-serum DMEM, 2μg/ml puromycin and 50 ng/mL recombinant human TGFβ2 (Peprotech, Rocky Hill, NJ), following an established protocol[53]. Cells were imaged using phase-contract light microscopy at 3, 7, and 21 days (d). The culture medium was changed every 3 days. Imaging experiments were repeated a minimum of 3 times. All experiments described above were concurrently carried out in regular mouse MSCs (C3H10T1/2; ATCC, Manassas, VA) but without puromycin, as these cells are not resistant to it.

*Western Blot Analysis (Supplemental Data)*

Cells were also collected for Western Blot (WB) immunoassays. However, due to limited sample availability, not all WB experiments were performed in triplicate, and have been included as supplementary data. Briefly, cells were collected for WB analysis in RIPA cell lysis buffer and HALT protease inhibitor (Invitrogen, Carlsbad, CA). Sodium dodecyl sulfate (SDS) was added at a 1:1 ratio and samples were prepared as described elsewhere[53]. Cell lysate collected from each well of the 12-well plate was run in its own lane (3 wells of each condition were run per individual experiment). Following protein separation, the gels were transferred to nitrocellulose membranes (Invitrogen), blocked in 5% milk in tris buffered saline (Boston Bioproducts, Ashland, MA) with 0.1% Tween20 (TBST) (Acros Organics, Morris Plains, NJ), and incubated overnight at 4 °C with Connexin-43, β-catenin, N-cadherin, and β-actin (loading control) primary antibodies raised in rabbit (all from Abcam, Cambridge, MA) in 5% bovine serum albumin (BSA) in TBST, at concentrations recommended by the manufacturer. The next day, blots were incubated with goat anti-rabbit HRP-linked secondary antibody (Invitrogen), developed using ECL chemiluminescence reagents (Invitrogen), imaged on a Genesys PXi (Syngene, Frederick, MD), and analyzed via band densitometry in ImageJ (NIH, Bethesda, MD), with all intensities normalized to their respective β-actin bands. As not all WB assays could be performed in triplicate due to limited sample availability, only qualitative results have been included as supplemental data to this study, to support the mechanical testing and imaging findings.

*Preliminary Analysis Suggests Disabling LINC via dnKASH inhibits tenogenesis in transfected MSCs (Supplemental)*

Transfected MSCs and controls were cultured in regular and tenogenic (+TGFβ2) media for up to 21 days to evaluate the tenogenic potential of dnKASH cells. Transfected control (Tr-control) cells did not receive TGFβ2, while Tr+TGFβ2 groups had 50 ng/mL TGFβ2 added to the media. All Tr-cells were supplemented with puromycin to select only for cells successfully transfected with the plasmid. Phase-contrast images (Supplemental Figure 3A-L) and Western Blot assays (Supplemental Figure 3M) showed changes in overall levels of several tenogenesis-associated proteins. We assessed only the 3-day timepoint due to inconsistencies in transfected cell viability at later timepoints. In contrast to published work[53] and vehicle controls showing an immediate decrease in N-cadherin levels during TGFβ2-induced tenogenesis, dnKASH cells maintained N-cadherin levels (Supplemental Figure 3J-L). Notably, transfected cells (controls and TGFβ2-supplemented) appeared to produce higher levels of β-catenin and Connexin-43 than vehicle control TGFβ2 non-transfected cells (Supplemental Figure G-L). As all these data are preliminary and based on n=2 or 3 Western Blot assays per protein, densitometry was not completed. Data are presented as supplementary evidence supporting a relationship between β-catenin, LINC, nuclear mechanosensing, and tenogenesis that will be assessed in future studies.
